# On the topochronic map of the human brain dynamics

**DOI:** 10.1101/2021.07.01.447872

**Authors:** P. Sorrentino, S. Petkoski, M. Sparaco, E. Troisi Lopez, E. Signoriello, F. Baselice, S. Bonavita, M.A. Pirozzi, M. Quarantelli, G. Sorrentino, V. Jirsa

## Abstract

Two structurally connected brain regions are more likely to interact, with the lengths of the structural bundles, their widths, myelination, and the topology of the structural connectome influencing the timing of the interactions. We introduce an in vivo approach for measuring functional delays across the whole brain using magneto/electroencephalography and integrating them with the structural bundles. The resulting topochronic map of the functional delays/velocities shows that larger bundles have faster velocities. We estimated the topochronic map in multiple sclerosis patients, who have damaged myelin sheaths, and controls, demonstrating greater delays in patients across the network and that structurally lesioned tracks were slowed down more than unaffected ones. We provide a novel framework for estimating functional transmission delays in vivo at the single-subject and single-fiber level.

**One-Sentence Summary:** A non-invasive estimation of the individual deterministic spatio-temporal scaffold underlying the evolution of brain dynamics.

## Main text

The localization approach, based on the observation that specific cognitive functions can be linked to the integrity of individual brain areas (e.g., Broca’s area in expressive aphasia) has yielded significant advancements in neurology over the past two centuries. However, higher-level cognitive functions could not be mapped to any specific area, suggesting that they might be inherently emergent and only exist as a result of the regulated interaction of multiple brain areas (*1*). Hence, network theory has been used as a framework to describe such large-scale patterns of interactions within the brain (*2*). With regard to structural connections, small-worldness (*3*) and scale-freeness (*4*) have been shown to characterize the organization of white matter connections. From the functional standpoint, most studies were based on assumptions of stationarity and focused on slow-evolving activity (typically in the timescale of seconds). The presence of time-averaged spatial patterns of correlated and anti-correlated activity was consistently reported (*5*). To some extent, these patterns could be inferred from the structural topology (*6, 7*). However, brain activity is far from stationary and instead displays fast, aperiodic, yet highly coordinated large-scale activities nested within the slower ones (*8*). What elements are needed to generate the “healthy” patterns of activations is not fully understood. Structural topology imposes constraints on the evolution of brain dynamics, as shown by the fact that the probability of two brain regions activating sequentially is proportional to the coupling intensity along the brain track linking them (*9*). However, structural topology alone is necessary, but not sufficient, to generate the observed large-scale patterns of activations. Delays, and perhaps noise, appear to also shape large-scale organization (*10*–*12*). The time it takes two gray matter regions to sequentially activate depends on multiple factors, such as the distance separating them, the properties of the white matter bundle (structural tract) linking them (such as its diameter and myelination), and the overall topology of the network embedding them (*13*). For this reason, estimating a comprehensive map of the delays poses great challenges, and most modeling studies assume homogeneous velocities of conduction (typically ranging from 2 to 10 meter/second) (*14*, *15*). Hence, the delays are conceptualized as purely dependent on distance (*11*, *12*, *16*). This is a necessary, yet suboptimal simplification, since we know (e.g. from routine neurological exams such as visual evoked potentials) that the velocities in highly myelinated, thick tracts can reach up to 150 m/sec (*13*). Combining magnetoencephalography (MEG) and tractography showed that it is possible to track the beginning and the spread of a regional perturbation (*9*). Building on this framework, we hypothesized that the time it takes a signal to go from one region to another might be a candidate proxy for the delay of transmission across any given structural tract. Shorter delays might relate, among other factors, to diameter and myelination. This should result in perturbations spreading with velocities that are not constant but, rather, rise as a function of the length of the white-matter tracts (given that longer tracts are typically also the thicker and more myelinated ones). Conversely, we expected the width of the distribution of the delays to be much narrower than what would be expected from the length of the white matter bundles alone (since the longer tracts are hypothesized to be the fastest). We further reasoned that, if our measurements are related to myelination, then lesions in the myelin sheath should increase the delays. To test this hypothesis, we used source-deconstructed MEG signals to estimate the delays, defined as the average time it takes a perturbation generated in region *i* to spread to region *j*, and then estimated the lengths of the white matter bundles in a cohort of 18 patients affected by multiple sclerosis and 20 healthy subjects using tractography. We used the delay in the subsequent recruitment of brain regions as a proxy for the conduction delay of an impulse between regions and estimated the corresponding velocities by dividing the tract lengths by the corresponding delays. First, we explored the distributions of the delays and velocities in healthy humans. Then, we evaluated the effect of the damage to the myelin sheath on the delays. To further verify the whole workflow, we applied the same procedure to a publicly available dataset based on combined electroencephalogram (EEG) recordings and tractography, and we were able to extract subject-specific delays and, thus, velocities, which had statistics that were similar to those obtained via MEG. The findings were further verified on an independent MEG/DTI dataset, using a different parcellation and a different source-reconstruction algorithm.

## RESULTS

### Delay estimation

In this study, we non-invasively estimated the functional delays in transmission across the network of white matter bundles in vivo. To this end, we combined source-reconstructed magnetoencephalography and tractography. Specifically, this framework relied on previous work showing that fast, structured bursts of coordinated activations (“avalanches”) are present in the human brain and that such activations spread preferentially along the white-matter bundles. To track avalanches, we *z*-scored each source-reconstructed MEG signal and defined the start of an avalanche as the moment when at least one region is “active,” i.e., when it is above threshold, and the end of an avalanche as the moment when no region remains active (Fig.1, A). Figure 1, B shows the main statistical properties displayed by neuronal avalanches. See supplementary material, Figure 1 – 3 for an extensive, subject-specific description. Once a region became active, the other regions were considered to be recruited in the avalanche if they went above the threshold in subsequent timesteps. The time it took for region *j* to be recruited by a perturbation that started in region *i* was then measured. Using this measurement, we built a matrix of delays in which rows and columns represented brain regions and each entry represented the delay between those two regions averaged across all the avalanches measured for one subject. Furthermore, using tractography, we also built a structural matrix, in which rows and columns are regions and entries are the (average) length of the tracts linking them. Finally, we obtained a velocity matrix, dividing the tract lengths by the corresponding delays. An overview of the pipeline is shown in Figure 2. The upper row of Figure 3 shows the average structural matrix for the controls (i.e., the tract lengths) with the corresponding distributions of the tract lengths on the right. The middle and bottom rows of Figure 3 respectively show the matrices and distributions of the delays and the velocities. Importantly, the structure of the delay matrix is related to the length of the structural tracts. However, the width of the distribution of the delays is much narrower than what would be expected given constant propagation velocities (see below). When estimating the velocities, we found that a consistent, fat-tailed distribution emerged. As expected, due to the heterogeneity of the tracts, the velocities appeared to be far from homogeneous, ranging from ~2 to ~60 m/sec. This range appears to be in accordance with known velocities of major myelin tracts (*13, 17, 18*). When plotting the distribution of the fastest edges (see glass brains in Figure 3), it appears that these were not evenly distributed across the brain and that the cross-hemispheric edges were preferentially selected as the fastest ones, a finding which agrees with the literature (*19*). We found that the delays grew as a function of the length of the tracts, as shown by the Spearman correlation between edge length and delays (*r* = 0.45, *p* = 3.85e-205). This relationship held at the individual level (Figure 4, panel B). However, while the delays were related to the tract lengths, they were not only determined by the lengths. In fact, the tract lengths ranged across an order of magnitude, whereas the corresponding increase in the delays was only moderate. For comparison, in Figure 4, panel C, we show the expected delays given constant velocities and reveal that the observed delays were remarkably steady despite the difference in the lengths of the structural tracts.

**Figure 1.**
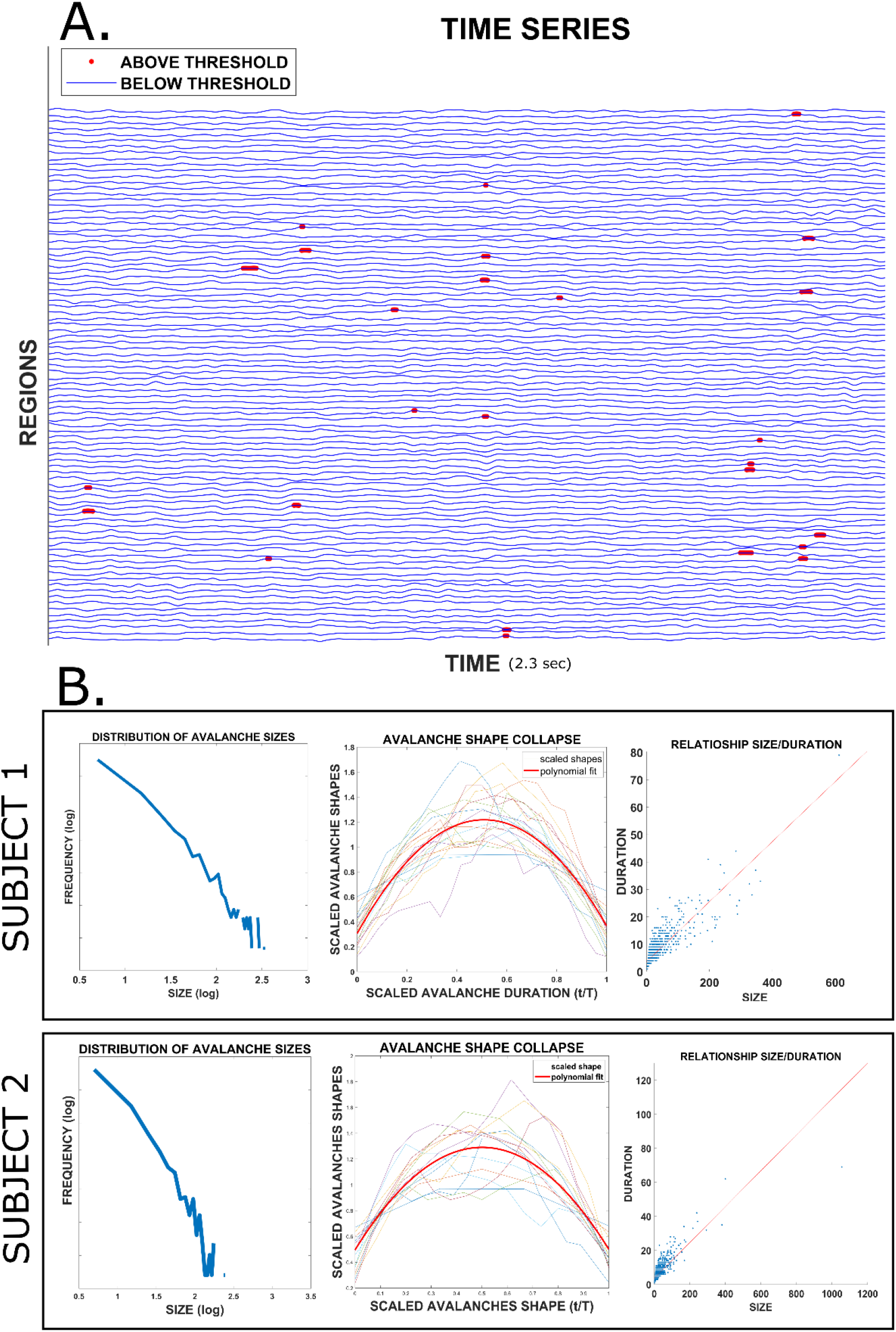
**Panel A.** Source-reconstructed MEG time-series. The time points that are below a threshold are represented in blue, while the timepoints above the threshold are in red (the red dots have been magnified). **Panel B.** Overview of the main statistical properties of neuronal avalanches for two randomly selected subjects: to the left, the distribution of the size of the avalanches (shown in log-log scale); in the middle, the “shape collapse”, i.e. the normalization of all avalanches to a unitary duration; to the right, the linear relationship between the size of the avalanches (i.e. the number of active bins) and their duration in time.

**Figure 2.**
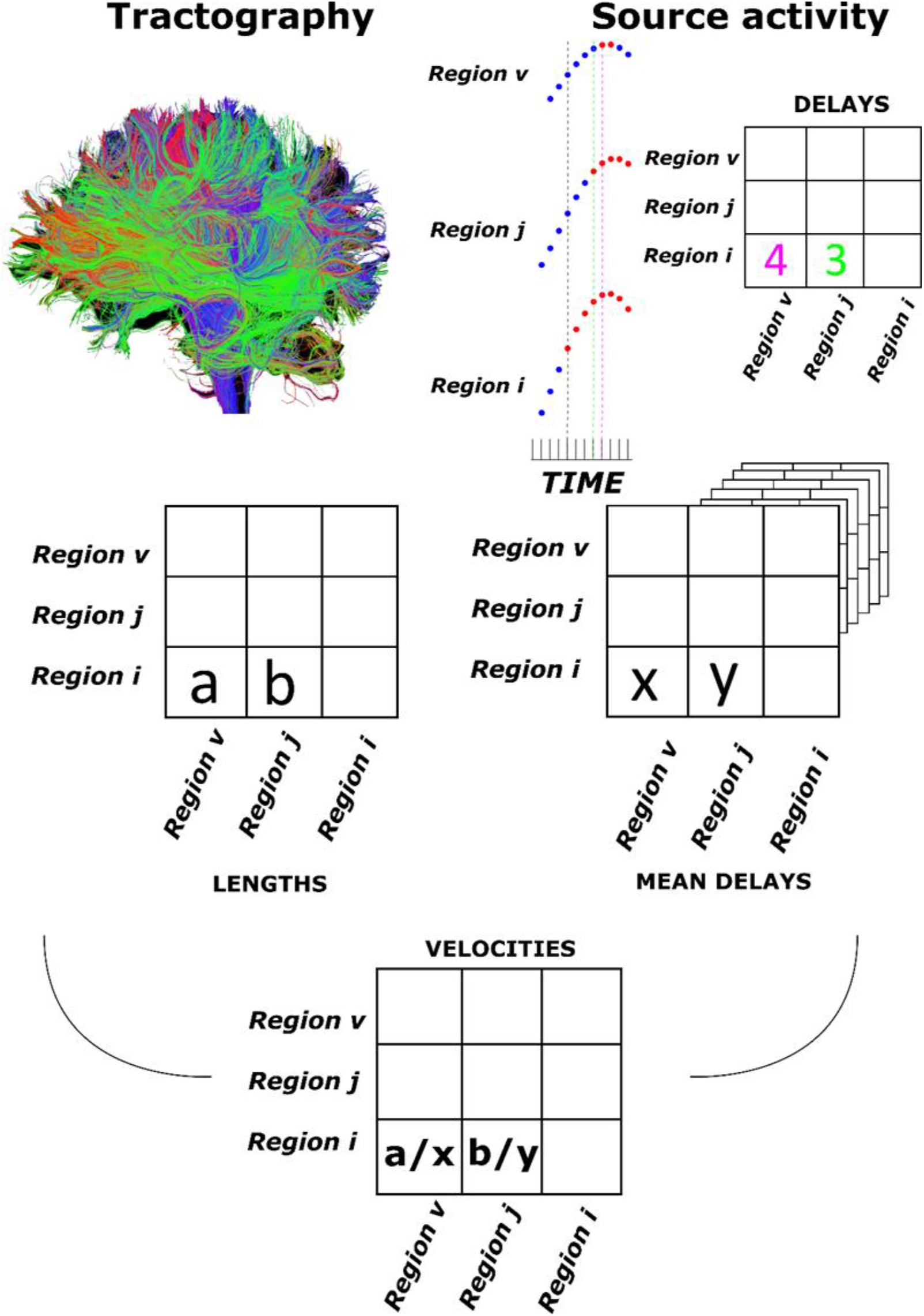
**Top left:** The structural connectome. **Middle left:** From the tractography, we obtained an estimate of each tract length. Averaging across subjects provided a group-level estimate. **Top right:** Avalanche-specific delay estimation procedure. The *z*-scored, source-reconstructed time series of three brain regions are represented. Time points are in blue if they are below the threshold (in this case, *z* = ±3), in red otherwise. After region *i* rose above the threshold (and, hence, the neuronal avalanche had started), it took region *j* three timesteps to be recruited and four timesteps for region z. The entries of the matrix are expressed in samples. Multiplying the number of samples by the sampling frequency allowed us to express the delays in terms of time. **Middle right:** Once avalanche-specific delay matrices were obtained, they were averaged across all avalanches belonging to one subject and then across subjects to obtain a group-specific matrix. **Bottom:** Each subject-specific length matrix was divided element-wise by the corresponding estimated delay to obtain a subject-specific delay matrix. Finally, the subject-specific velocity matrices were averaged to obtain a group-specific velocity matrix.

**Figure 3.**
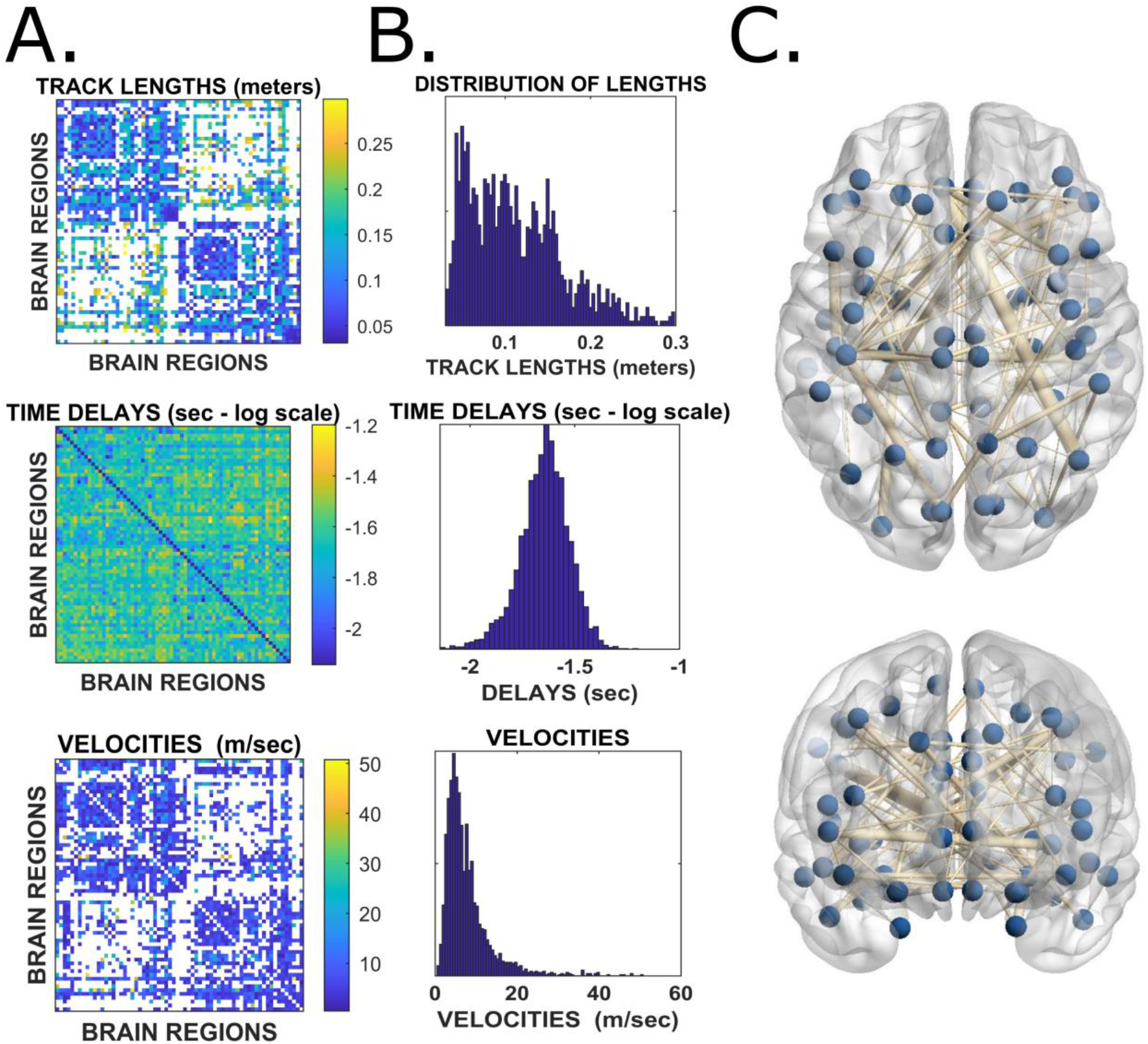
**A.** Top: Track lengths. Rows and columns represent brain regions, and the color code conveys the corresponding length of the track linking any two regions. Middle: Group average of the delays. Rows and columns represent brain regions, while the color code represents the average time it took region *j* to become active provided region *i* had been active earlier. Values are in seconds, and reported on a log-scale to highlight the texture of the delay matrix, which appears to be strongly correlated to the length matrix. Bottom: As before, brain regions are represented as rows and columns, while the matrix entries represent the velocities, expressed in meters/second. **B.** Top, middle, and bottom are the histograms corresponding to the matrices for the lengths, delays, and velocities, respectively. **C.** The glass-brain provides an overview on the topography of the functional edges with fast functional velocities. In particular, the blue dots represent brain regions, and the edges signify the delays. Only edges with a minimum velocity of 15 m/sec are shown. Given this lower-bound limit, the width of the edge is proportional to the velocity of the edge.

**Figure 4.**
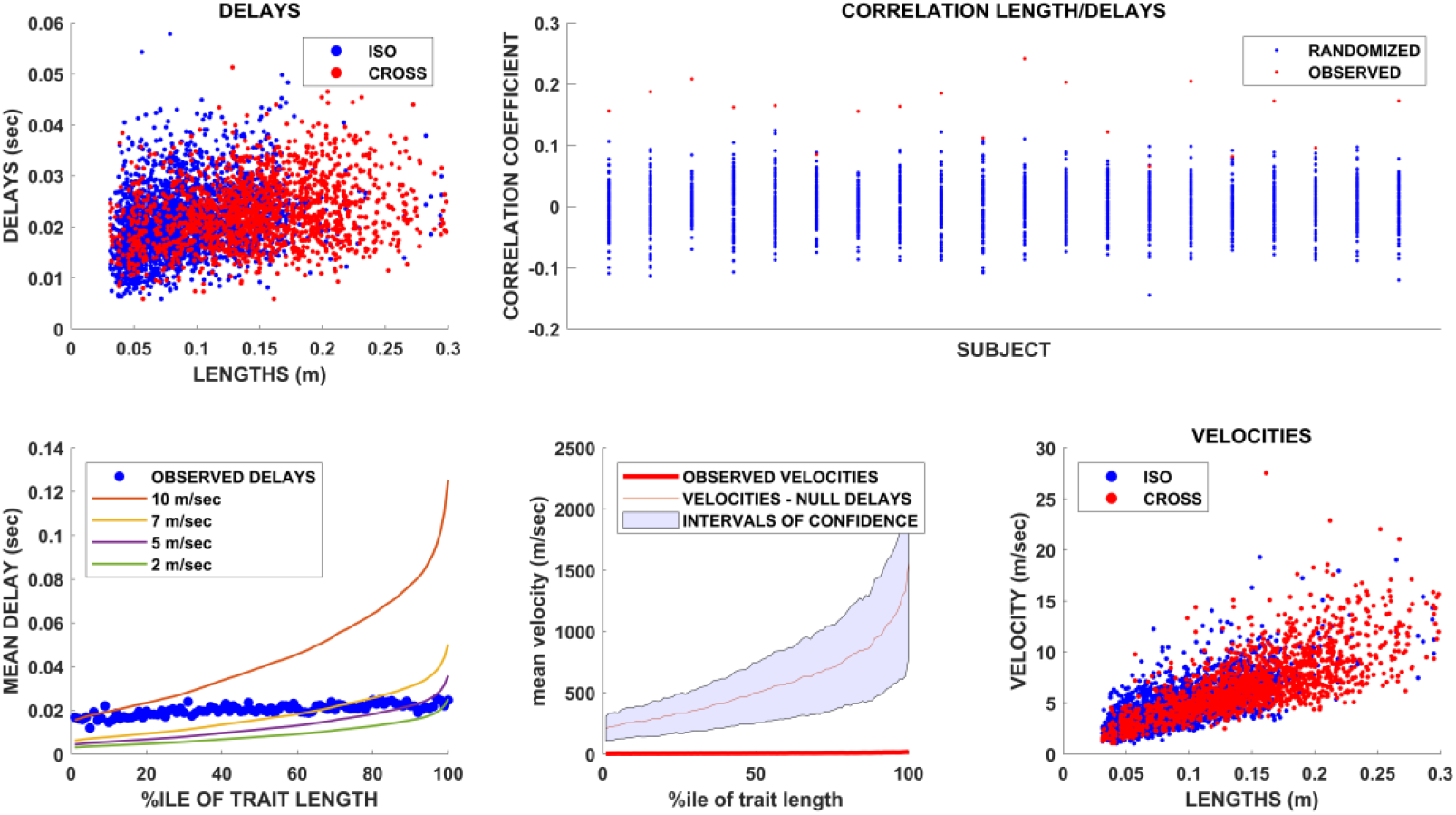
**Top left:** Group–level relationship between track lengths and functional delays. Blue dots represent iso-hemispheric edges, while red dots represent cross-hemispheric edges. **Top right:** For each of the 20 healthy controls, the red dot represents the intensity of the relationship between the tract lengths and delays, calculated using Spearman’s correlation coefficient. For each subject, the avalanches were randomized by shuffling the time sequence while preserving the spatial pattern of the active regions within each timestep. Based on these newly randomized avalanches, the edgewise average delay was calculated and then related to the track lengths. The procedure was repeated 1000 times per subject. The resulting Spearman’s correlations are represented as blue dots. **Bottom left:** For each subject, the delays were averaged according to the percentile of the corresponding length. Hence, each blue dot represents the average delay across all edges whose length belonged to the (patient-specific) *n*^th^ percentile. The colored lines show the delays that would be expected if the velocities were homogeneous across all edges. In this case, the growth of the delays followed the growth of the track lengths. **Bottom middle:** The red dots represent the observed velocities (averaged according to the percentile of the track length. The red line represents the average velocity estimated based on the delays derived from the randomly shuffled avalanches. **Bottom right:** Group-level relationship between the velocities. Blue dots represent iso-hemispheric edges, while red dots represent cross-hemispheric edges.

### Statistical validation

Next, we sought to establish whether our results were likely to have been obtained by chance alone. To this end, we built a surrogate dataset based on randomized avalanches in which we shuffled the time sequence while preserving the spatial structure. In other words, for each avalanche, the time points were randomly shuffled, but the regions recruited at any given time point were fixed. By doing this, the time-structure of the recruitment of regions was disrupted, but the purely spatial component was retained. After the permutations, the average delays were again computed for each edge. The procedure was repeated 100 times, yielding a hundred surrogate delays for each edge. First, we compared the delays retrieved from the random timeseries to the tract lengths, obtaining 100 Spearman’s *r*s, to which we compared the observed correlation, *p* < .001. We then used the random surrogates to compute random velocities, i.e., dividing each edge length by the corresponding delay derived from the random surrogates. Using the same method, this procedure was carried out 100 times (once for each of the 100 random surrogates), retrieving a distribution of the correlations obtained between tract lengths and random velocities (i.e., the velocities computed using random delays). As expected, the random velocities appeared to be more strongly related to the track lengths than the observed velocities (*p* < .001). In other words, when we divided the lengths by the randomized delays, the resulting velocities became a function of the track length alone. On the other hand, when we divided the track lengths by the observed delays, the longer tracts appeared to be faster than if the delays were only a function of distance. The mean (red line) and upper and lower bounds (shaded area) of the surrogate velocities derived from surrogate delays (grouped by the percentile of the corresponding track length) are shown in Figure 3, D. The delays derived from the surrogate data led to a higher estimate of the velocities compared with the observed ones. Finally, as shown in panel E, we confirmed, using real data, that the transmission velocities grew as a function of the length of the tracts, such that the longer tracts were also the faster ones. All in all, this part of the analysis showed a finely regulated relationship between the delays and the lengths of the structural tracts, which implies non-homogeneous functional velocities.

### Delay estimation in multiple sclerosis patients

Next, we sought to test our framework in patients affected by multiple sclerosis, which is a prototypical disease in which myelin in the central nervous system is selectively attacked by the immune system (*20*). We excluded patients with severe functional impairment (Expanded Disability Status Scale, EDSS < 7) so that we could obtain a picture that is likely to be influenced more by demyelination than by degenerative phenomena (*21*). We expected to observe greater delays (and lower velocities) in the patient population compared with the controls. As shown on the left side of Figure 4, the average delay per percentile was consistently higher in the multiple sclerosis patients compared with the controls. The empirical cumulative distribution function confirmed the difference in the delays in patients compared with the controls (KS test, *p* < .001). Note that, in this analysis, similar to what we did before, we grouped the delays according to percentile across the whole group. Hence, the average value refers to the average delay of all the edges that fall within a given length percentile, with the percentile defined for each subject. We then moved on to an edgewise comparison. To this end, for each patient, the average delay of each edge was compared to the average delay of each corresponding edge in the controls. Hence, we obtained the difference between the edge-specific delay and the corresponding average delay in the controls for each patient. To base our findings on a more stable estimate, we considered only edges for which a delay estimate was available for each of the 20 controls. Figure 4C shows the distribution of the differences between the delays (patients minus controls). The distribution is not centered around zero, as would be expected if the patients did not have longer delays. Instead, the distribution is heavily skewed toward positive values, implying longer delays in the patients compared to controls. Finally, we investigated edge-specific lesions. In fact, while it is reasonable to expect globally greater delays in the patients compared with the controls, the delays corresponding to edges that were lesioned might be more lengthened compared with the delays corresponding to healthy edges. For each patient, we classified edges as healthy or lesioned, based on the presence or absence of structural damage. Under the null hypothesis that the delays corresponding to structurally lesioned edges do not differ from the delays corresponding to non-lesioned edges, we calculated the average delay difference in random samples of non-lesioned edges (in patients), with the size of the sample equal to the number of lesioned edges. We repeated this procedure 1000 times and compared the obtained distribution to the observed difference in the lesioned edges. The results are shown in Figure 5, panel D. The delays in the lesioned edges slowed more (with respect to the corresponding delays in the healthy population) than the healthy edges. Hence, selecting a subset of edges based on the structural information, we retrieved a difference in the temporal structure of the functional dynamic. This difference would not be expected if the structural damage was unrelated to the delays, as shown by the permutation analysis (*p* < .001).

**Figure 5.**
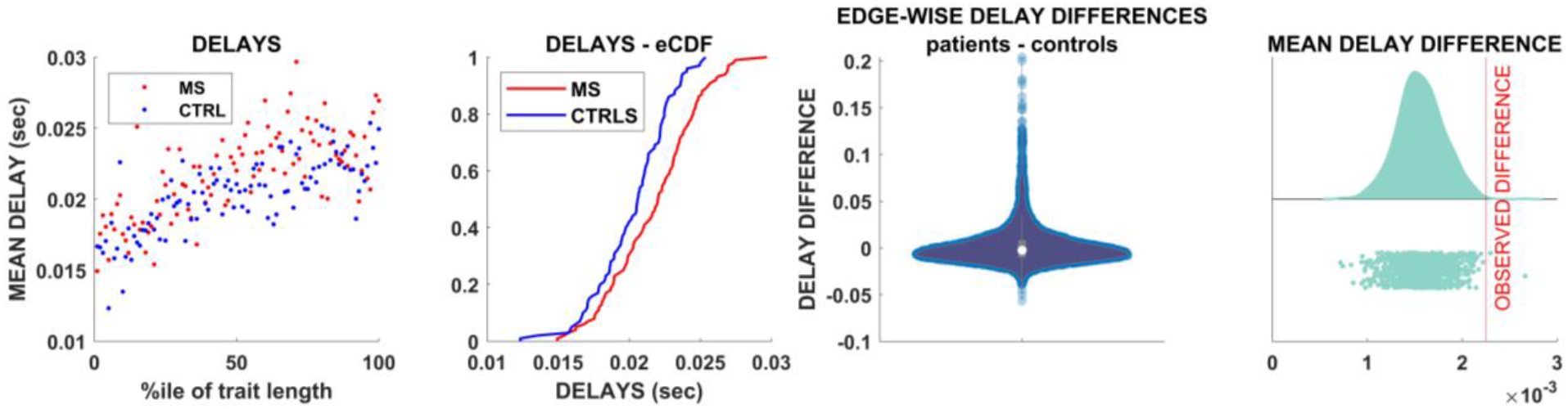
**Panel A.** Each dot represents the average delay for each percentile of track length. Blue dots represent controls, while red dots represent patients. **Panel B.** Empirical cumulative distributions of the delays. The blue line corresponds to the average delays of the controls, while the red lines correspond to the average delays of the MS population. **Panel C.** The violin plot shows the edgewise differences between delays in each patient and the average delay across all the controls for the corresponding edge. **Panel D.** The vertical red line marks the observed average edgewise difference in the delays calculated only for delays that were lesioned. The distribution to the left shows the average delay difference observed after selecting a random sample based on a thousand randomizations, of edgewise delay differences (i.e., ignoring the information about the structural integrity) with the size of each random sample equal to the number of lesioned edges. Th results enabled us to reject the null hypothesis that edge-specific lesions would not affect the delays.

### Replication in independent datasets

The results of the delay estimation were tested using an independent dataset based on co-registered MRI and EEG, and all the main findings were confirmed (see Supplementary Material Figure 1). Note that the cleaning and source-reconstruction algorithms were different in this dataset as compared to ours, showing robustness to both the technique used and data processing. Furthermore, we replicated the main results using different parcellations (AAL and DKT), varying the *z*-score threshold (2.5 and 3.5 in addition to the 3.0 reported here), varying the binning (see methods), and using yet another source-reconstruction algorithm (i.e. the “residual variance” method). The residual variance is a particular dipole-fit approach that involves the minimization of the signal that remains unexplained by a current source model in which dipoles are assumed to have fixed position and fixed or varying orientation. Conversely to LCMV, there is no linear constrain involved in the minimization process (*22*). Furthermore, a larger MEG/tractography dataset, involving 47 young healthy subjects, was also used to further explore the robustness of our findings. These results were replicated using the AAL atlas. Finally, we varied the minimum size of the avalanches used to compute the delays, from taking all avalanches into account, up to selecting only those longer than 10/15 samples. These analyses are reported in the Supplementary Material, Figures 4–13). Finally, we confirmed that the delay matrices showed convergence as the sample size increased, as a further check on the validity of our results (Supplementary Material, Figure 14).

## DISCUSSION

### The topochronic map

In this study, we developed a novel method to quickly and non-invasively obtain an estimate of the time delays that occur when locally generated perturbations spread to other brain regions. Our methodology was able to capture the presence of homogeneous delays across brain regions, even though the length of the structural tracts grew according to a fat-tail distribution (*23, 24*). If the delays were only dependent on the track lengths, they should be longer for longer tracts (i.e., it should take longer to cover a greater distance). This effect should be major, given that longer white-matter tracts are roughly one order of magnitude longer than shorter ones (*2*). The fact that the delays do not scale with increased track lengths implies one or more compensatory mechanisms. Such mechanisms might involve varying axonal diameters, myelination, and network effects. In particular, myelination greatly affects the velocity of propagation (*17*, *18*, *25*). In fact, given our estimated delays, we found that, in the healthy subjects, the velocities ranged from ~3 to ~60 m/sec. This observation is in sharp contrast with the typical simplifying assumptions, made in modeling studies, that velocities are constant, which would be expected to lead to a broad range of delays across the network. The fact that the transmission between the regions connected by the longest edges is faster than those with shorter edges is likely important in terms of the unfolding of the dynamics and should not be surprising considering the experimental results on stimulation although until this current study solid measurements of personalized links on the whole-brain level had been missing. Hence, non-invasive estimates of the temporal constraints in vivo are highly relevant for modeling individual dynamics. Furthermore, many of the fast edges connect regions that are highly central to the brain network, thus, likely causing reverberation that greatly and nonlinearly impact the average transit time (*26*). Although we can only speculate, we think that it is possible that homogeneous delays, that is, having perturbations reach wide-spread brain regions or arrive at a focal point from wide-spread regions simultaneously, would be favorable for allowing the brain to have simultaneous access to information from multiple locations across the brain, a concept which fits well into the framework of global workspace theory (*27*). This supposition finds support in that white matter damage impairs conscious access in multiple sclerosis patients (*28*, *29*). A reduction in the complexity of the spatio-temporal spreading of such perturbations (referred to as “neuronal avalanches” within the framework of critical dynamics) has been shown to be related to states of reduced consciousness (*30*) as well as to neurodegeneration (*31*, *32*). Importantly, our analyses focused on rare, intermittent, large-scale bursts of activations, which have been consistently observed in human brains (*33*). The importance of such rapid transients to large-scale brain dynamics is confirmed by recent findings showing that the patterns of functional connectivity are shaped by specific, short moments in time (*34*). Furthermore, the fact that avalanches preferentially spread along structural tracts indicates that functional delays might be used as a proxy to estimate the velocities across individual tracts (*9*). As we investigated the topographic distribution of the fast edges, we found that they were spatially distributed non-homogeneously. Importantly, fast connections seem to be preferentially cross-hemispheric, a finding that is not surprising from a neuroanatomical standpoint, provided that these connections are mediated, for example, via the corpus callosum (*35*). While our data was based on broad-band data, the patterns that emerged were not dominated by the occipital alpha frequency, as it is often the case with M/EEG data. However, we wish to stress that such results might have been biased by the tractography, which may have preferentially estimated tracts in specific anatomical regions (*19*). Hence, further validation is needed to confirm this finding, which should be considered explorative. The replication of our results in an independent MEG dataset as well as in a publicly available multimodal EEG-MRI dataset (*36*) indicates the reliability of our findings by increasing the probability that they are not modality-specific.

### Implications in neurological diseases and multiple sclerosis

As explained, myelination is believed to greatly influence conduction velocity and, hence, to modulate delays. To test this, we measured the delays in multiple sclerosis patients. As expected, the delays were greater in multiple sclerosis patients than in matched healthy controls. Selecting MS patients at a fairly early stage should make the role of demyelinating lesions more prominent than the role of degeneration (*37*). However, this was not quantified and remains a potential source of confounds that needs further investigation. To provide edge-specific information about the time delays, we used subject-specific lesion masks to separate lesioned edges from non-lesioned ones in the patients. We focused the analysis on the delays to avoid potential biases induced by unreliable length estimates due to the demyelinating lesions (*38*). We observed, for both the lesioned and non-lesioned edges, that functional delays increased in the multiple sclerosis patients, in accordance with the hypothesis that damage to the myelin would provoke longer delays. This also provides support for the claim that our measurements are related to temporal patterns of delays that are imposed on the overall structural connectome. The fact that avalanches propagate more slowly along lesioned edges shows the relevance of the direct pathways in determining the edge-specific delays given the expected relationship between the structural integrity of the track and the velocity of propagation along it. However, it is important to stress that the role of network effects cannot be easily disentangled (*2*). In fact, the transmission in patients was slower even across unaffected tracts. This might be interpreted as an expression of the fact that the delays likely depend from a combination of both direct and indirect paths through which a perturbation can potentially travel between two regions. In this sense, one does not expect two regions that are linked by a healthy edge that is embedded in a diseased network to communicate as quickly as two regions that are also linked by a healthy edge as well as embedded in a healthy network. Other contributing factors may include the erroneous classification of damaged but sub-threshold edges as healthy.

### Cross-dataset validation and limitations

The strength of this study is that the track lengths and the delays were estimated using two different techniques, making it unlikely that the relationship is spurious or tautological. We tested our results by changing both the binning parameter (see Methods) and the *z*-score threshold to define the spreading of the perturbations and the brain parcellation, again showing the robustness of our findings. As similar results were obtained using EEG, utilizing a different preprocessing pipeline, applying a different algorithm to source-reconstruct, and performing a further parcellation, our confidence in these results is further strengthened (see Supplementary Materials). The fact that the relationship between length and velocity was maintained at the subject level is remarkable. One limitation that should be considered, however, is the fact that we used the DKT and the AAL atlases, both of which are coarse grained. However, finer grained parcellations, while optimal for structural MRI, would have been below the resolution for MEG and, hence, might have created spurious results.

### Conclusions

We propose that, beyond the topology of functional connections imposed by the spatial scaffolding (*9*), the conjugate property of connectivity is temporal in nature and complementary to the structural topology (*39*). Together, spatial and temporal constraints reveal the topochronic framework from which oscillatory brain activity emerges. Applying tools from statistical mechanics and dynamical system theory to “reverse engineer” the pattern of delays that occur in the living brain, we non-invasively measured the large-scale patterns of functional delays that occurs in the human brain at rest. The pattern of delays is basically similar across individuals. Hence, the time it takes an (internally generated) local impulse to affect other regions is not only a function of the track length but is also heavily modulated by the properties of the track itself and, globally, by the large-scale structure of the network. Including subject-specific delays provides the potential to improve virtual, personalized in-silico brain models (*40*). Importantly, the fact that delay estimation can be obtained from EEG, a widely available technique, allows nearly every facility to include subject-specific delays into personalized brain models. Furthermore, large cohorts will be needed to get normative data from large, stratified cohorts, to create priors to be included in personalized models, in case subject-specific estimations are not available. In conclusion, we proposed a simple method for combining multimodal imaging within the framework of statistical mechanics, to derive subject-specific topochronic maps of large-scale brain dynamics in both healthy and diseased populations.

## Data and materials availability

Code will be available on GitHub. The MEG data and the reconstructed avalanches are available upon request to the corresponding author (Pierpaolo Sorrentino), conditional on appropriate ethics approval at the local site. The availability of the data was not previously included in the ethical approval, and therefore data cannot be shared directly. In case data are requested, the corresponding author will request an amendment to the local ethical committee. Conditional to approval, the data will be made available.

## MATERIALS AND METHODS

### PARTICIPANTS

The participants were recruited at the outpatient clinic of the Institute for Diagnosis and Cure Hermitage Capodimonte (Naples, Italy). The diagnosis of multiple sclerosis was made according to the revised 2017 McDonald criteria (*41*). Exclusion criteria were age < 18 years, clinical relapse and/or steroid use in the 3 months before the study, inability to understand and complete “patient reported outcomes” and cognitive evaluation, or inability to undergo the MRI scan. All patients underwent a neurological clinical examination, Expanded Disability Status Scale (EDSS) scoring, the Symbol Digit Modalities Test (SDMT) to measure cognitive impairment, the Fatigue Severity Scale (FSS), and the Beck Depression Inventory (BDI). The controls for the MS cohort were selected from among the caregivers and spouses of the patients. Genetic relatives were not allowed as controls. The subjects for the second, larger healthy cohort were selected as described in (*9*). The independent EEG/DTI dataset is described in (*42*). The demographics and main clinical and radiological features of the MS cohort are summarized in Table 1. The study was approved by the local Ethics Committee (Prot.n.93C.E./Reg. n.14-17OSS).

**Table 1.** Features of the multiple sclerosis cohort

|  | **CTRL** | **MS** | *p*-value |
| --- | --- | --- | --- |
| **Age (years)** | 45.8 (± 11) | 44.9 (± 9.9) | .8 |
| **Education (years)** | 13.6 (± 3.8) | 13.8 (± 5) | .9 |
| **Gender (m/f)** | 6/14 | 6/12 | - |
| **Disease duration (months)** | - | 187.7 (± 131.8) | - |
| **EDSS** | - | 4.5 (± 1.9) | - |
| **SDMT** | - | 40.3 (± 13) | - |
| **FSS** | - | 36.1 (± 14) | - |
| **BDI** | - | 12.8 (± 1.3) | - |
| **LL** | - | 12959 (± 12253) | - |

#### MRI acquisition and processing

Each MRI scan was performed immediately after the MEG recording on the same MRI scanner (1.5 Tesla, Signa, GE Healthcare). Analyzed sequences included echo-planar imaging for DTI reconstruction (TR/TE 12,000/95.5 ms, voxel 0.94×0.94×2.5 mm^3^, 32 equally spaced diffusion-sensitizing directions, 5 B0 volumes) and 3D-FLAIR volume for WM lesion segmentation (TR/TE/TI 7000/145/1919 ms, echo train length 170, 212 sagittal partitions, voxel size 0.52×0.52×0.80 mm^3^). Preprocessing of the diffusion MRI data was carried out using the software modules provided in the FMRIB software library (FSL, http://fsl.fmrib.ox.ac.uk/fsl). All diffusion MRI datasets were corrected for head movements and eddy current distortions using the “eddy_correct” routine (*43*), rotating diffusion sensitizing gradient directions accordingly (*44*), and a brain mask was obtained from the B0 images using the Brain Extraction Tool routine (*45*). A diffusion-tensor model was fitted at each voxel, and fiber tracks were generated over the whole brain by deterministic tractography using the Diffusion Toolkit (*46*) (FACT propagation algorithm, angle threshold 45°, spline-filtered, masking by the FA maps thresholded at 0.2). Two cortical study-specific ROI datasets were obtained by masking the ROIs available in the AAL atlas (*47*) and in an MNI space-defined volumetric version of the Desikan-Killiany-Tourville (DKT) ROI atlas (*48*) using the GM tissue probability map available in SPM (thresholded at 0.2). This was done to limit the endpoints of the fibers to cortical and adjacent juxtacortical white matter voxels in the subsequent ROI-based analysis of the tractography data. The analysis was replicated twice to ascertain how robust the method was when used with specific brain parcellations. To obtain the corresponding patient-specific ROI sets, each participant’s FA volume was spatially normalized (*49*) to the FA template provided by FSL using SPM12, and the resulting normalization matrices were inverted and applied to the two ROI sets. Additionally, for each subject, the MS lesion map was obtained by segmenting the 3D-FLAIR volume using the lesion prediction algorithm (*50*) implemented in the Lesion Segmentation Tool (LST toolbox version 3.0.0, www.statistical-modelling.de/lst.html) for SPM. The 3D-FLAIR volume was then co-registered to the EPI of the patient (*51*), and the coregistration matrix was applied to the corresponding WM lesion volume, which was resampled by nearest-neighbour interpolation, thus obtaining the patients’ lesion masks, which would be coregistered with the DTI volume. Finally, for each patient the average length of the fibers connecting each pair of ROIs and the percentage of voxels crossed by those fibers that had MS lesions were calculated separately for the AAL and DKT ROI sets, using an in-house routine written in Interactive Data Language (IDL, Harris Geospatial Solutions, Inc., Broomfield, CO, USA).

#### MEG pre-processing

MEG pre-processing and source reconstruction were performed as in (*52*). Preprocessing and source reconstruction operations were carried using the Fieldtrip Toolbox (*53*). Each participant underwent a MEG recording, composed of both eyes-closed resting-state segments of 3’30” each. Four anatomical coils were applied on the head of each participant and their position was recorded along with the position of four head anatomical points, to identify the position of the head during the recording. Eye blinking (if present) and heart activity were recorded through electro-oculogram (EOG) and electrocardiogram (ECG), to identify physiological artifacts (*54*). An expert rater checked for noisy signals and removed them. An anti-alias filter was applied to the MEG signals, acquired at 1024 Hz, before being filtered with a fourth order Butterworth IIR band-pass filter (0.5-48 Hz). We used principal component analysis (*55*, *56*) and supervised independent component analysis (*57*) to remove the environmental noise and the physiological artifacts (recorded with EOG and ECG), respectively.

#### Source reconstruction

Signal time series were reconstructed using both the AAL and DKT atlases (*47*, *58*), which consist of 116 and 84 ROIs, respectively. The reconstruction took place utilizing the volume conduction model proposed by Nolte (*59*). Based on the native MRIs of each subject, the linearly constrained minimum variance (LCMN) (*60*) beamformer was applied to reconstruct the signal sources based on the centroids of each ROI (*60*). ROIs belonging to the cerebellum were excluded due to the low reliability of their source reconstruction (*61*), for a total of 90 ROIs in the AAL atlas and 84 ROIs in the DKT atlas.

#### Neuronal avalanches and branching parameter

To study the dynamics of brain activity, we based our analysis on “neuronal avalanches”. First, the time series for each ROI was downsampled to 512 Hz and discretized by calculating the *z*-score; then the positive and negative excursions beyond a threshold were identified. The main results reported here refer to a threshold equal to 3 standard deviations (|*z*| = 3), but thresholds of 2.5 and 3.5 were also tested. A neuronal avalanche began when at least one ROI went above the threshold (|*z|* >3) and ended when all the ROIs were below the threshold (*62*, *63*). Before proceeding with the analyses, we binned the data, to ensure that we captured any critical dynamics, if present. To estimate the suitable time bin length, for each subject, for each neuronal avalanche, and for each time bin duration, the branching parameter σ was estimated (*64*, *65*). In fact, systems operating at criticality typically display a branching ratio of ~1 (*66*). The branching ratio was calculated as the geometrically averaged (over all the time bins) ratio of the number of events (activations) between the subsequent time bin (descendants) and that in the current time bin (ancestors) and then averaging the avalanche-specific branching ratio over all the avalanches (*67*). More specifically:

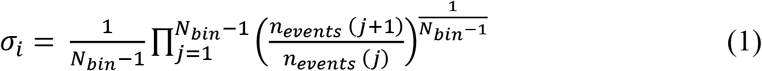

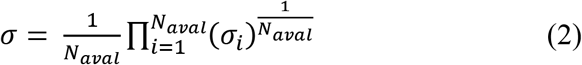

Where *σ_i_* is the branching parameter of the *i*^th^ avalanche in the dataset, *N_bin_* is the total number of bins in the *i*^th^ avalanche, *n_events_*(*j*) is the total number of events active in the *j*^th^ bin, and *N_aval_* is the total number of avalanches in the dataset. We tested bins from 1 to 5, and picked 3 for further analysis. However, the branching ratio was around 1 for all the bins we tested, and the results were unchanged for other bin durations (see Supplementary Material). The results shown were derived when taking into accounts avalanches longer than 5 time bins. However, we repeated the analysis taking into account avalanches longer than 10 and than 15 time bins, as well as taking all the avalanches into account, and the results were unchanged (see supplementary Materials, Figures 7 to 9.

#### Estimation of delay matrices

The delays were estimated for each avalanche. The procedure is schematized in Fig.1. In an avalanche, from the moment region *i* activated, we recorded how long it took region *j* to activate. These are what we considered to be delays. Hence, for each avalanche we obtained a matrix, in which the rows and columns represented brain regions and the entries contained the delays. We then averaged across all the avalanches belonging to one subject, obtaining an average *ij^th^* delay. The average was performed disregarding zero entries, since each avalanche-specific matrix is very sparse. With this procedure, a subject-specific delay matrix was built. Averaging across subjects (again discarding zero entries) yielded a group-specific matrix.

#### Statistical testing

To build null models to test the functional delays estimates, we randomized the temporal order of the avalanches without changing the spatial structure. To do this, the order of the time bins of each avalanche was randomly shuffled, while the pattern of active regions within each bin was kept unchanged. The delays were then estimated using the shuffled avalanches, and the avalanche-specific delays matrices were averaged across each subject and then across groups, as described previously. We compared the observed delays with the surrogate delays. We then used the delays estimated from the random surrogates to compute functional velocities (dividing the length of the structural edges by the surrogate delay). Finally, we computed the growth of the velocities as a function of the length of the structural tracks and compared this with the observed distribution. To test differences between the distribution of the delays in the healthy subjects and patients, we used the Kolmogorov-Smirnov test. To perform the edge-wise comparison of the delays in the healthy vs. lesioned edges in MS patients, we used permutation testing (*68*). In short, we tested the null-hypothesis that lesions in the edges would not have an impact on the delays. First, we calculated the average edge-wise difference between the delays in each patient and the average delay in the corresponding edge in the controls. Then, we randomly selected a subset of the differences in the delays with the same size as the number of lesioned edges and computed its average. We repeated this procedure 1000 times, building a distribution of the differences in the delays that are to be expected by randomly drawing a subsample of the edges. Finally, we compared this distribution to the observed difference in the lesioned edges to obtain the probability of observing the data under the hypothesis that the edge specific lesions would not slow down the functional transmission.

---

**Fig. S1.**
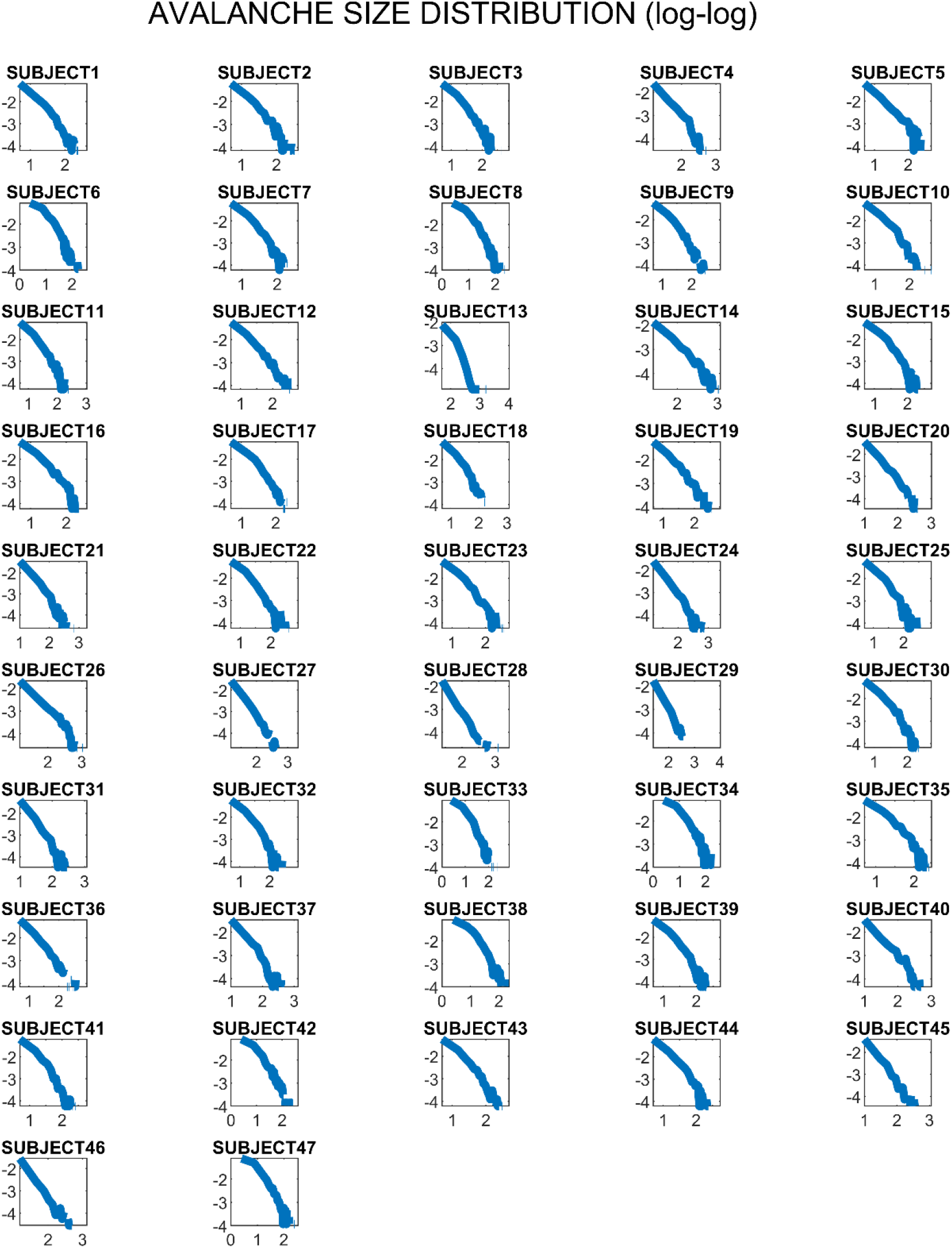
Subject-specific distribution of the size of the neuronal avalanches (this refers to the MEG replication dataset).

**Fig. S2.**
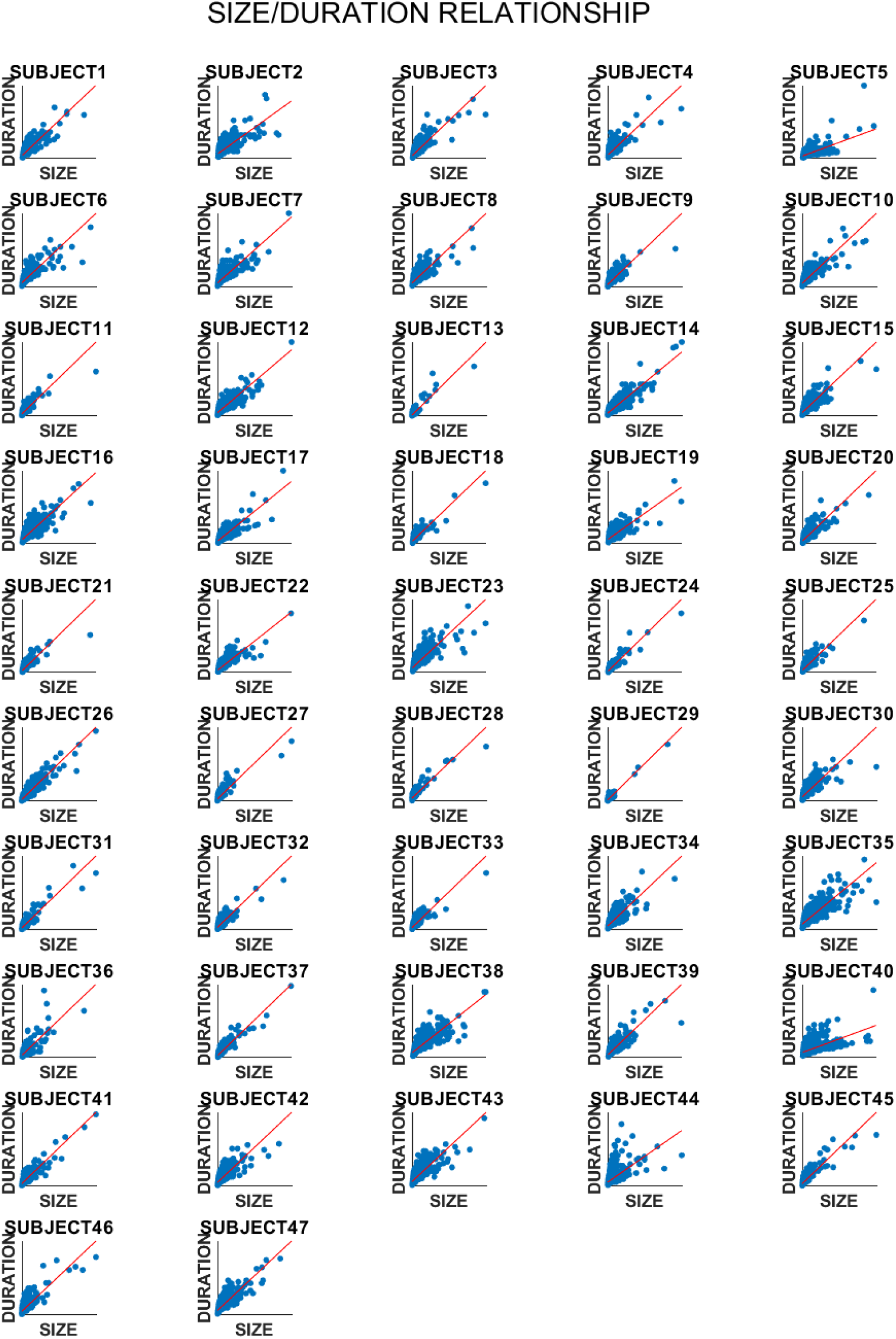
Subject-specific scatterplot showing the relationship between avalanche size and duration (this refers to the MEG replication dataset).

**Fig. S3.**
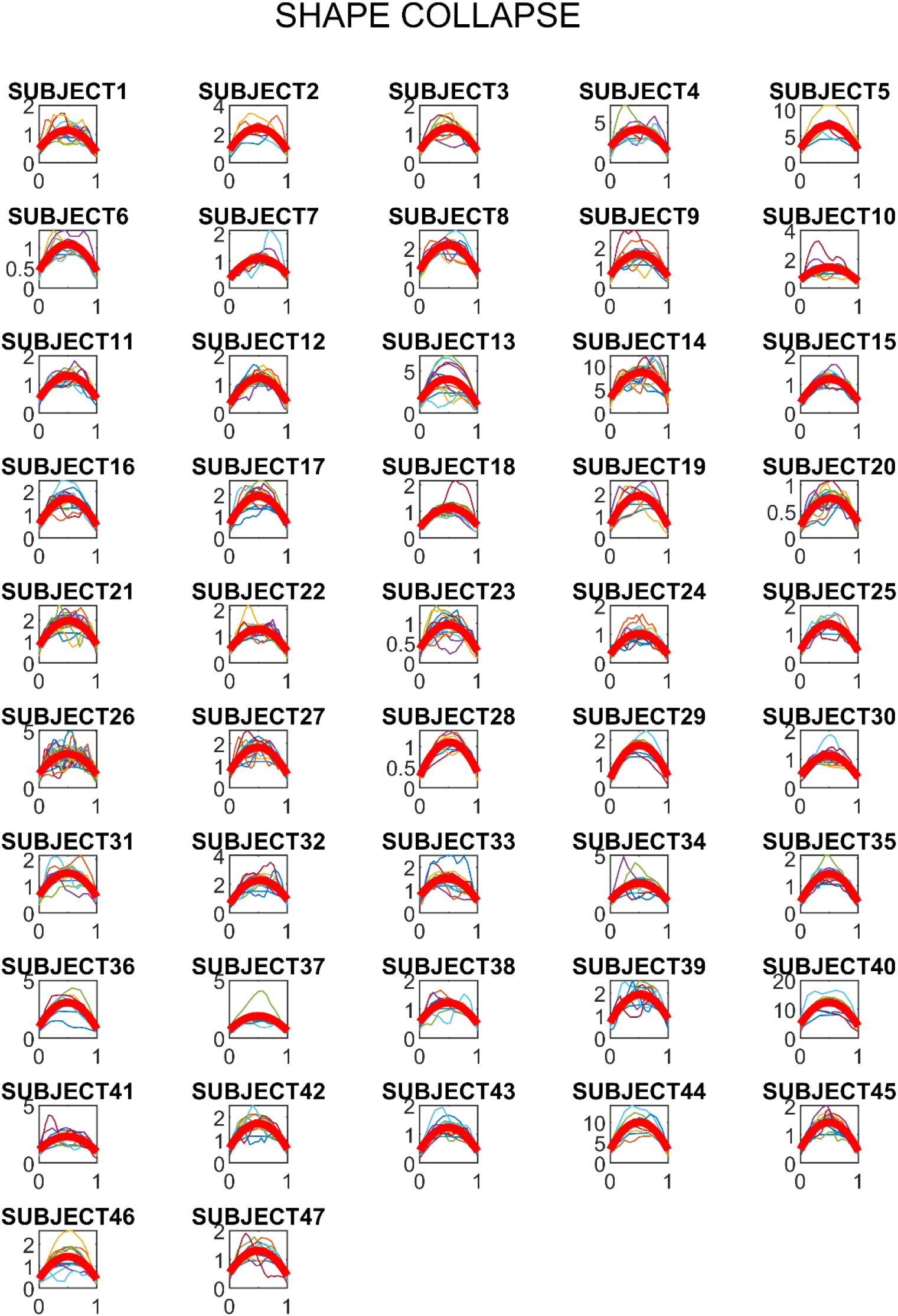
Subject-specific shape-collapse of the avalanches, with best polynomial fit shown in red (this refers to the MEG replication dataset).

**Fig. S4.**
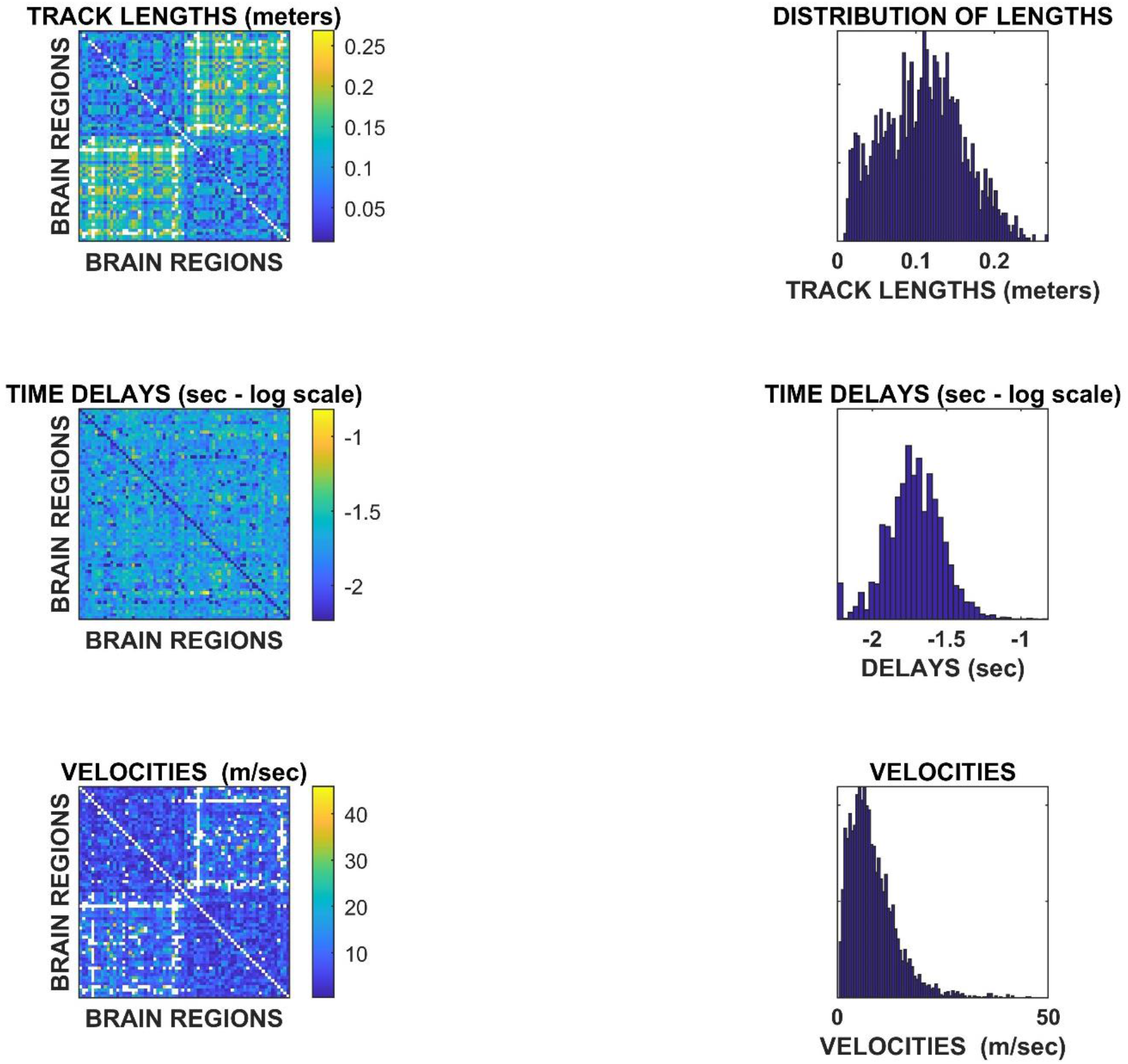
Replication of the main results on an independent cohort, based on source-reconstructed EEG and tractography.

**Fig. S5.**
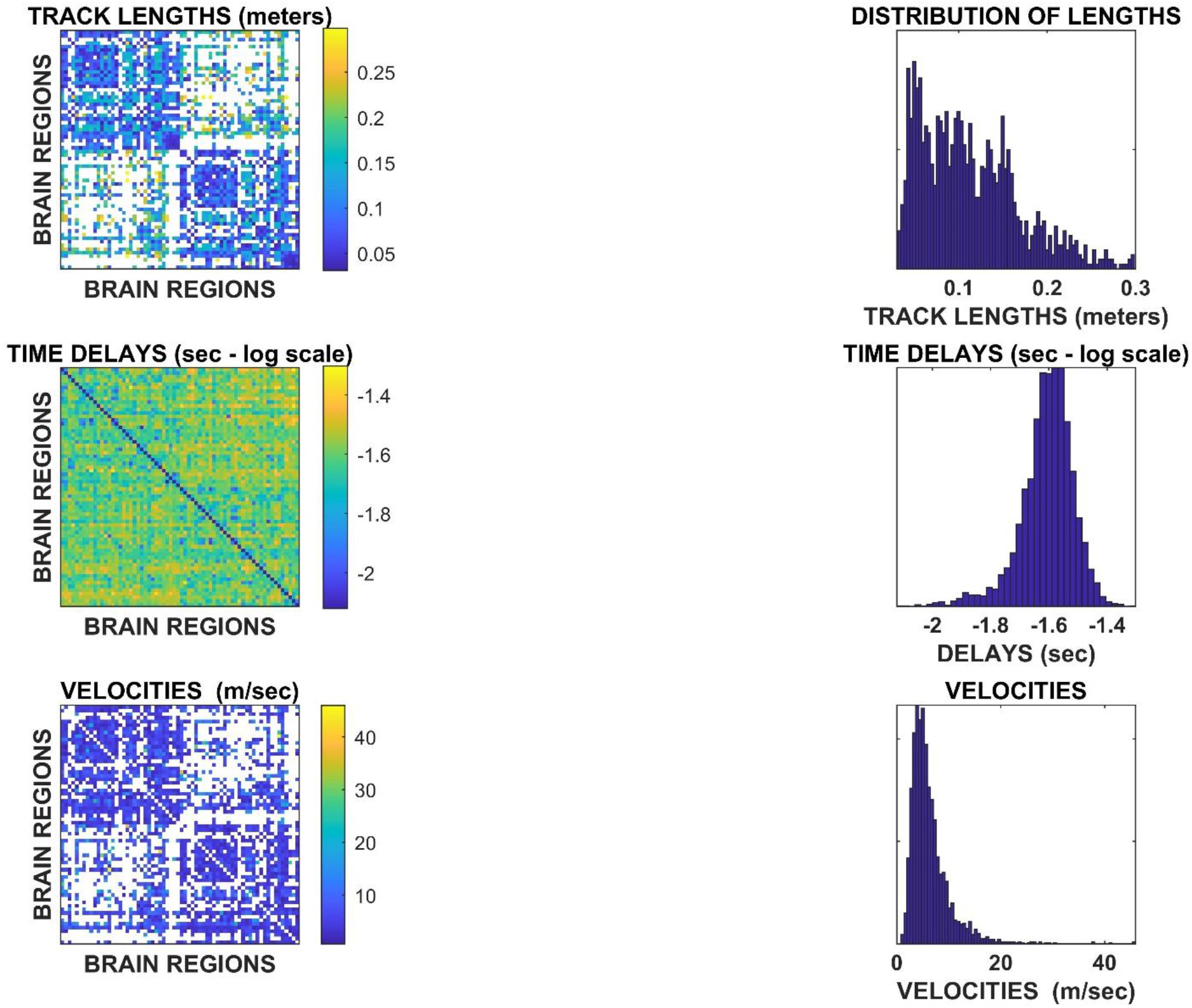
Replication of the main results with a *z*-score >±2.5.

**Fig. S6.**
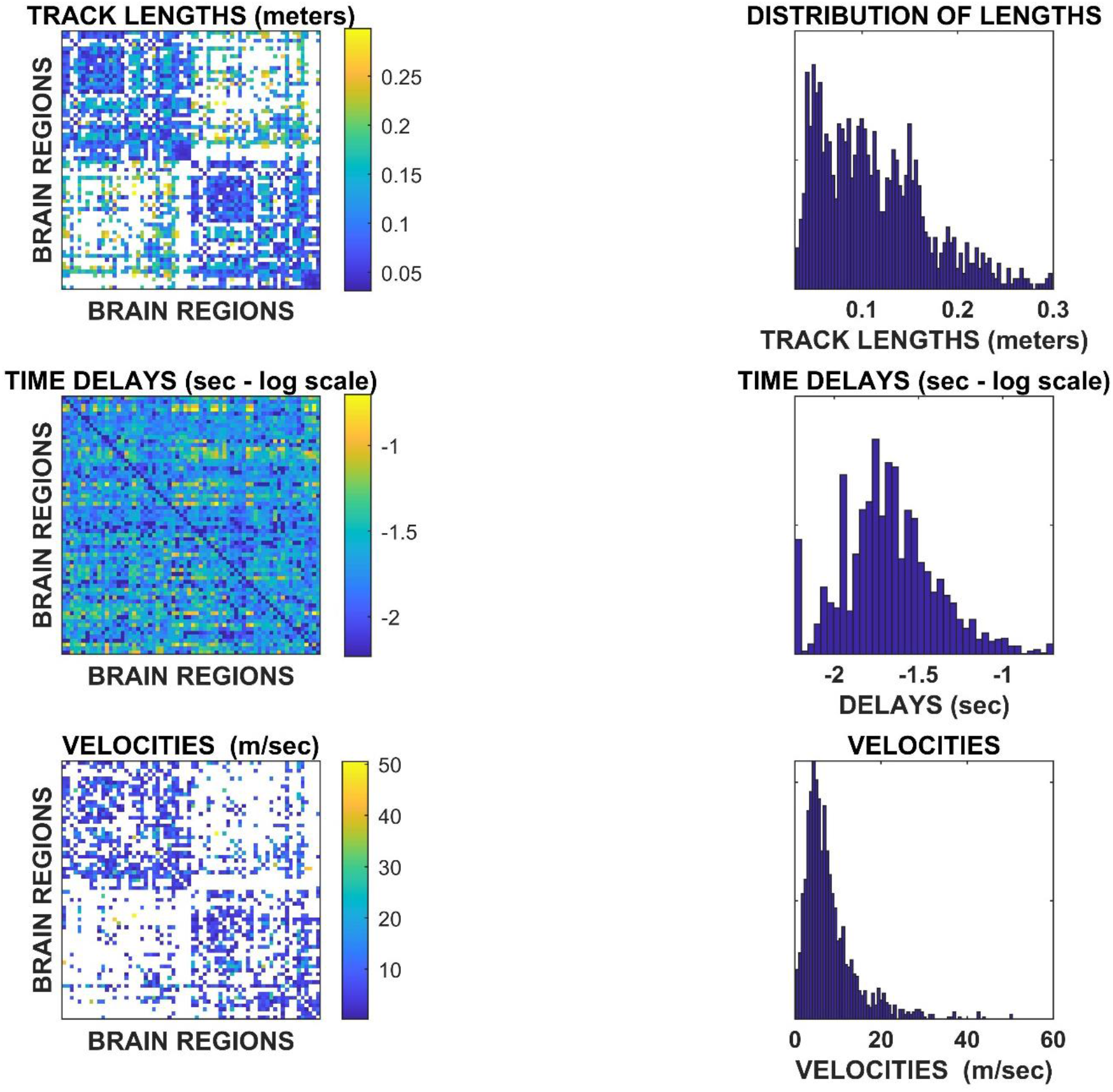
Replication of the main results with a *z*-score >±3.5.

**Fig. S7.**
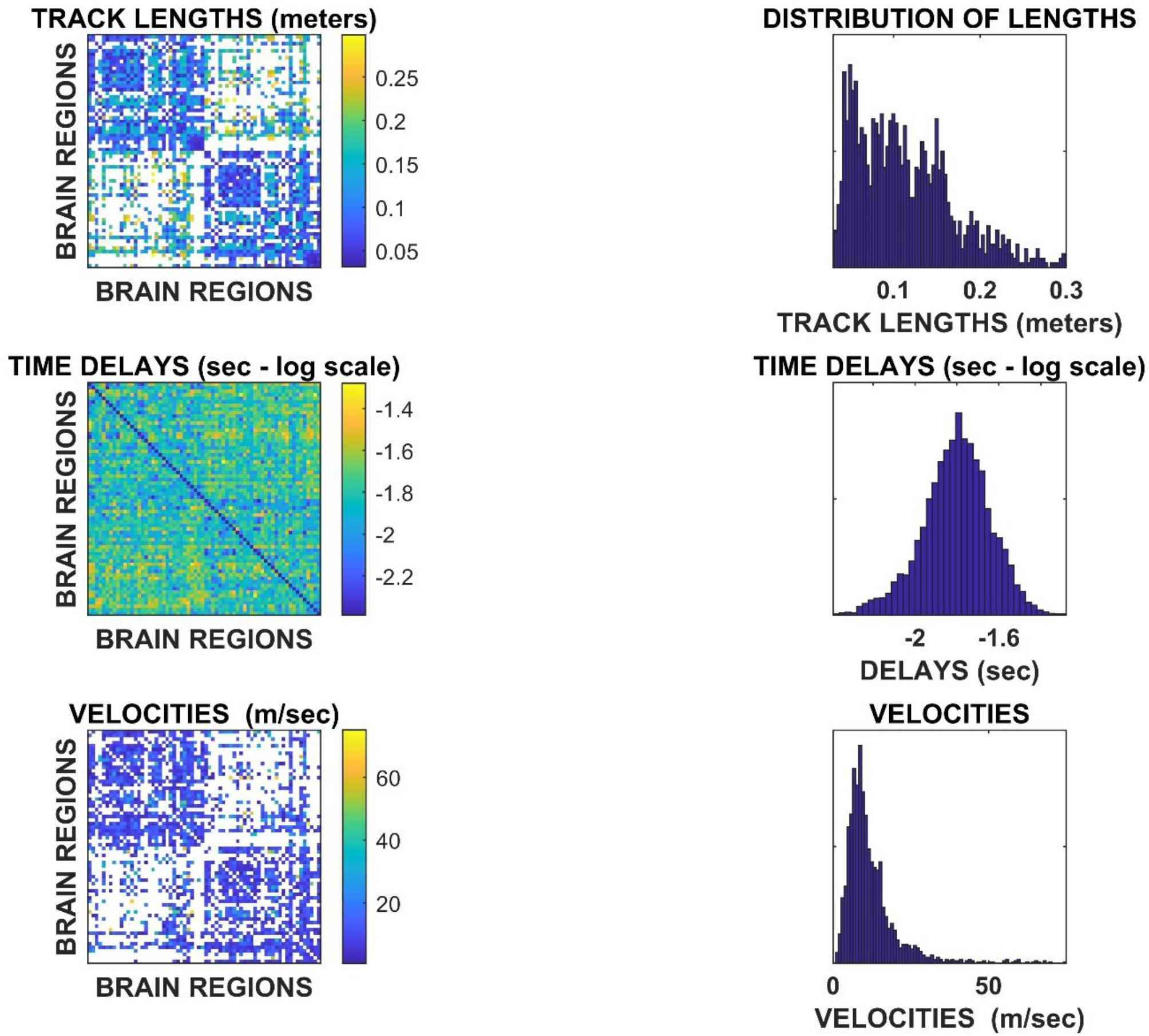
Replication of the main results with binning = 2.

**Fig. S8.**
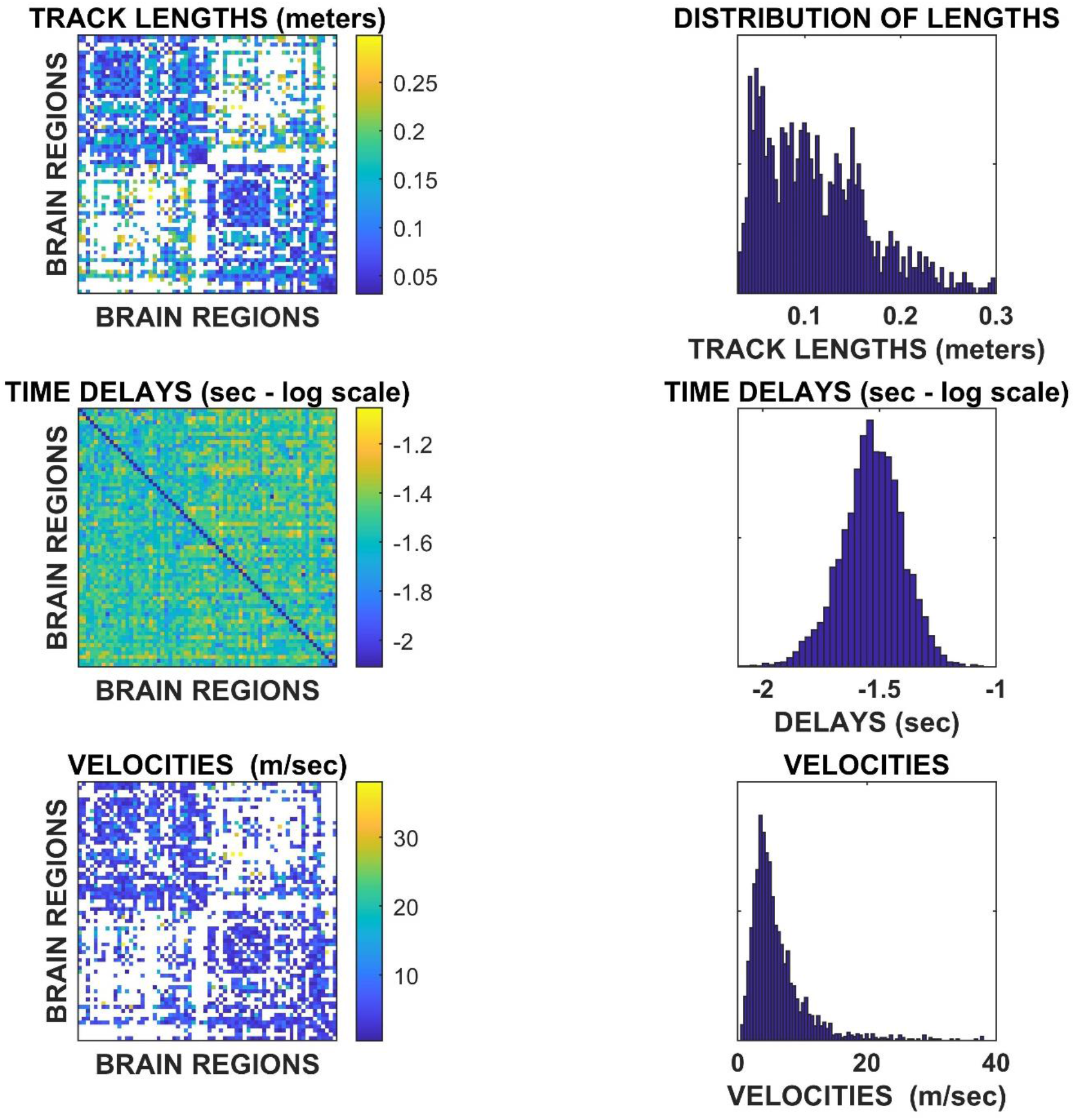
Replication of the main results with binning = 4.

**Fig.S9.**
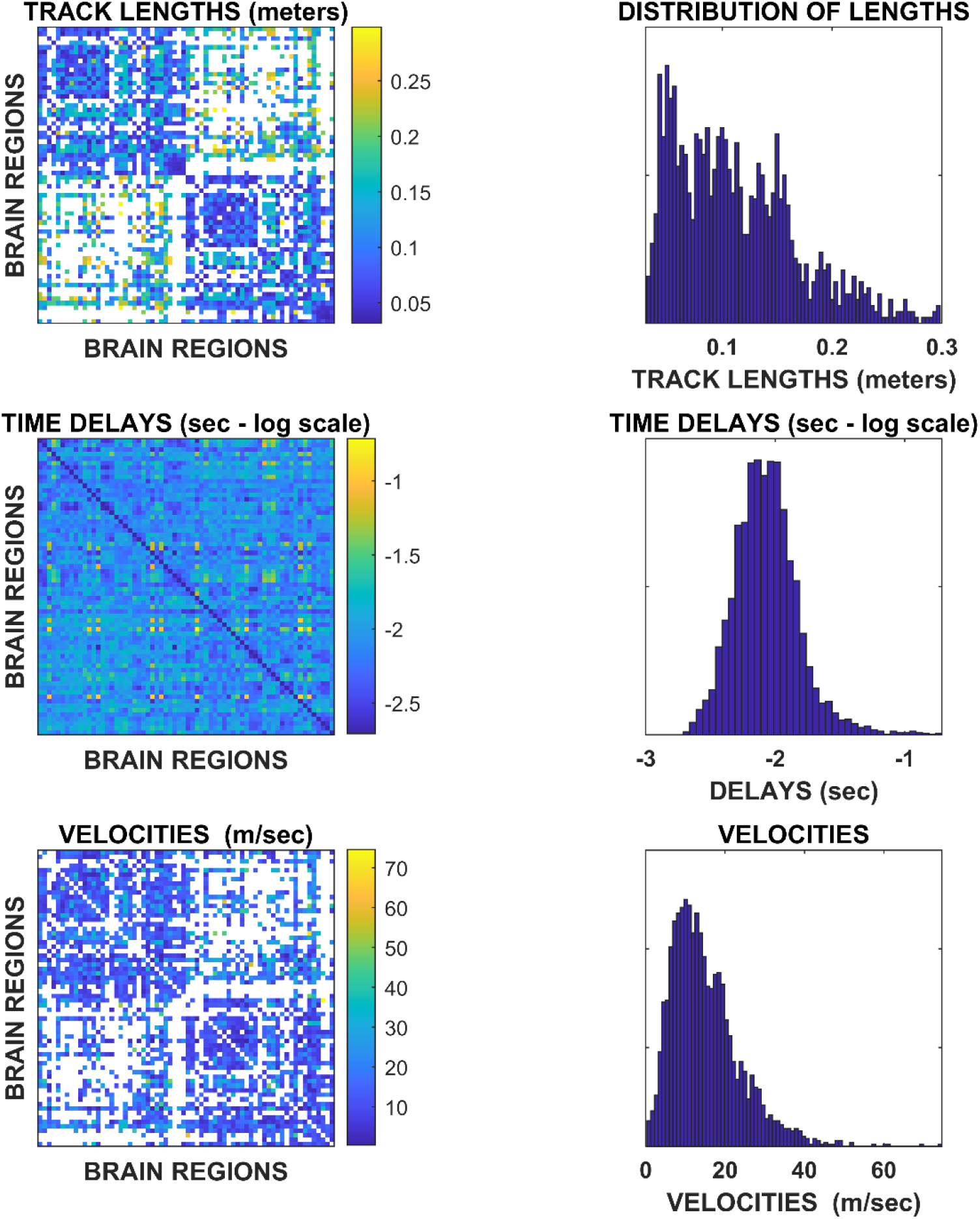
Replication of the main results using the “residual variance” as a source-reconstruction algorithm.

**Fig.S10.**
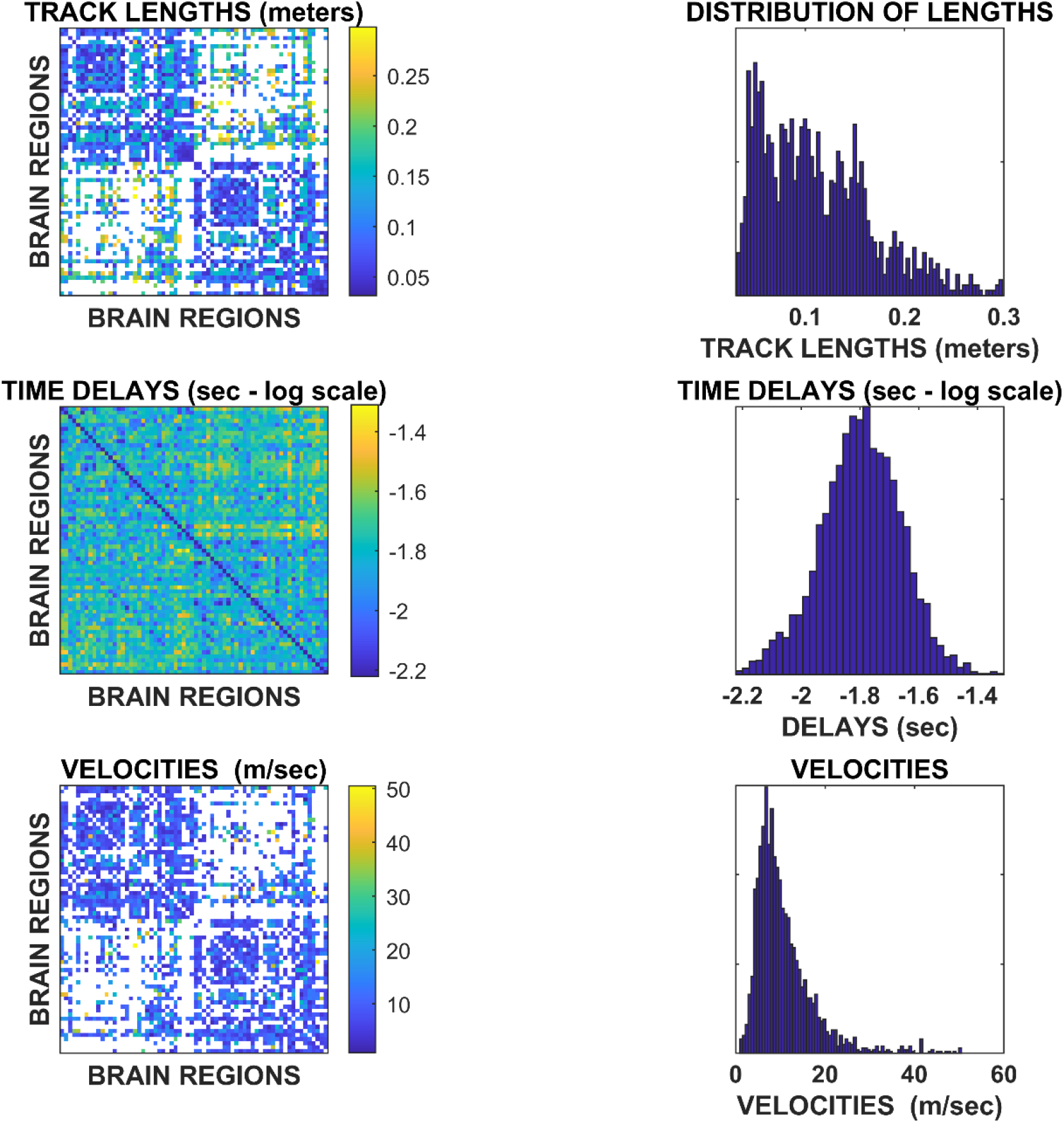
Replication of the main results using all available avalanches.

**Fig. S11.**
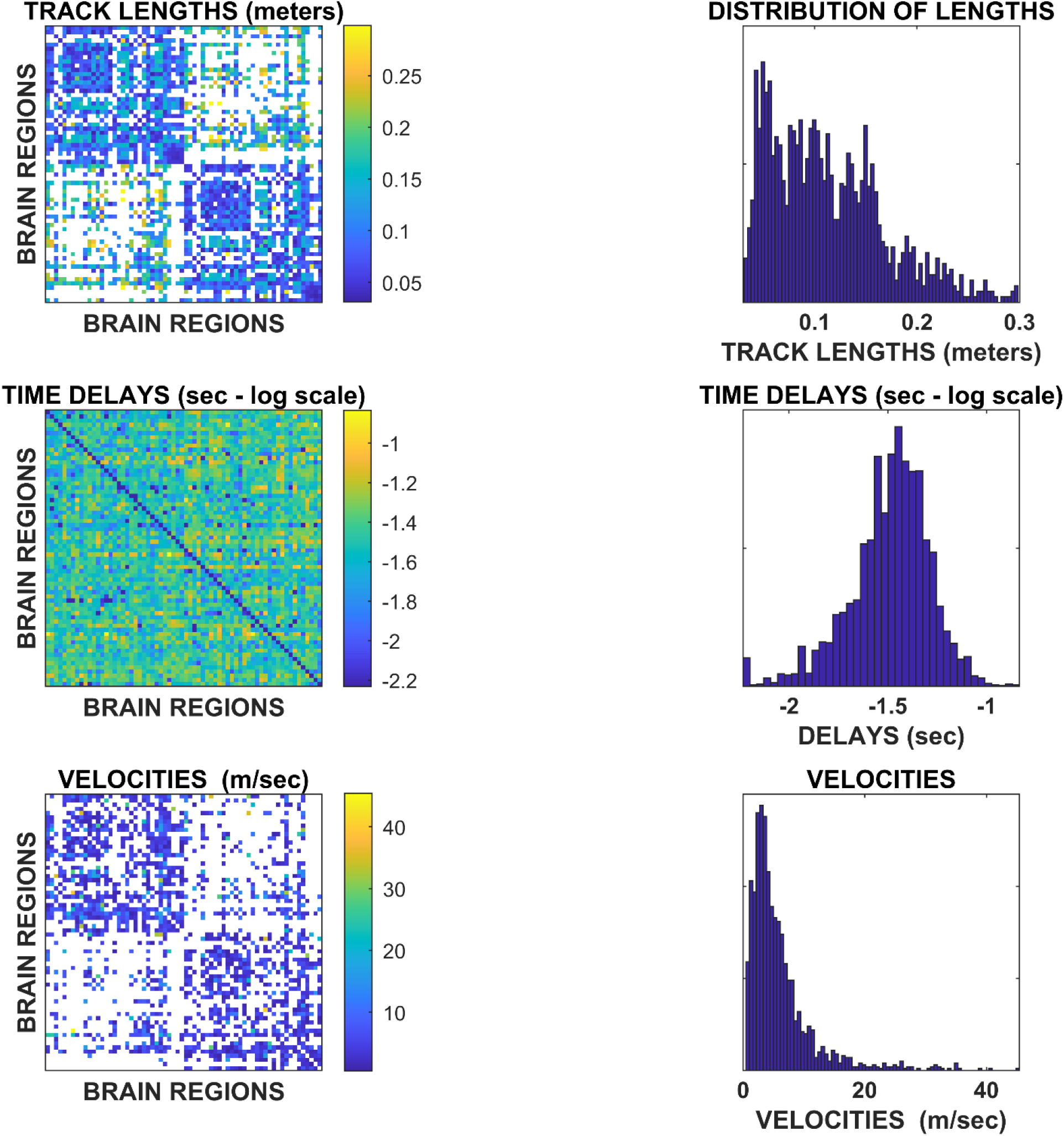
Replication of the main results using available with size greater than 10.

**Fig. S12.**
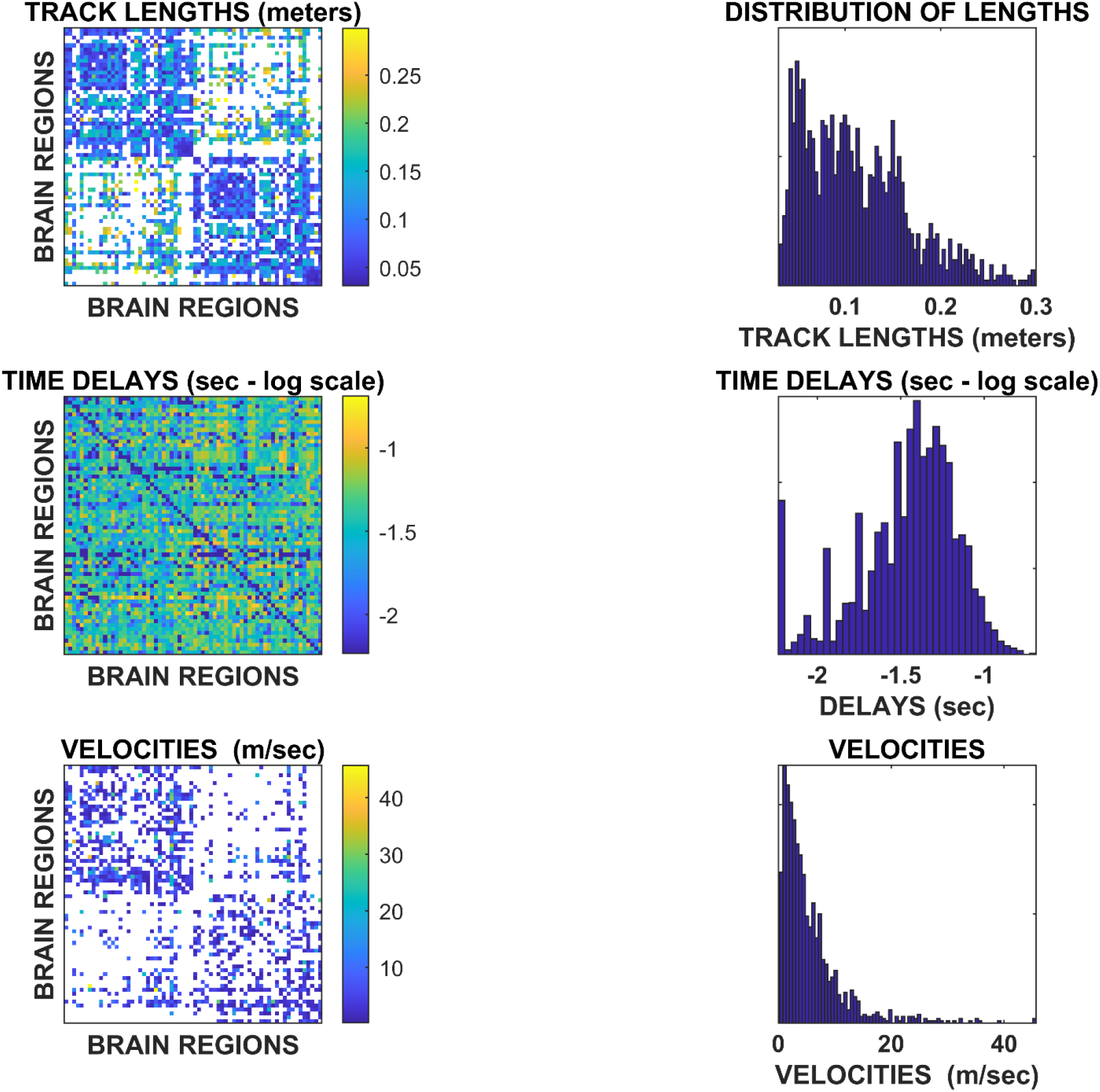
Replication of the main results using available with size greater than 15.

**Fig. S13.**
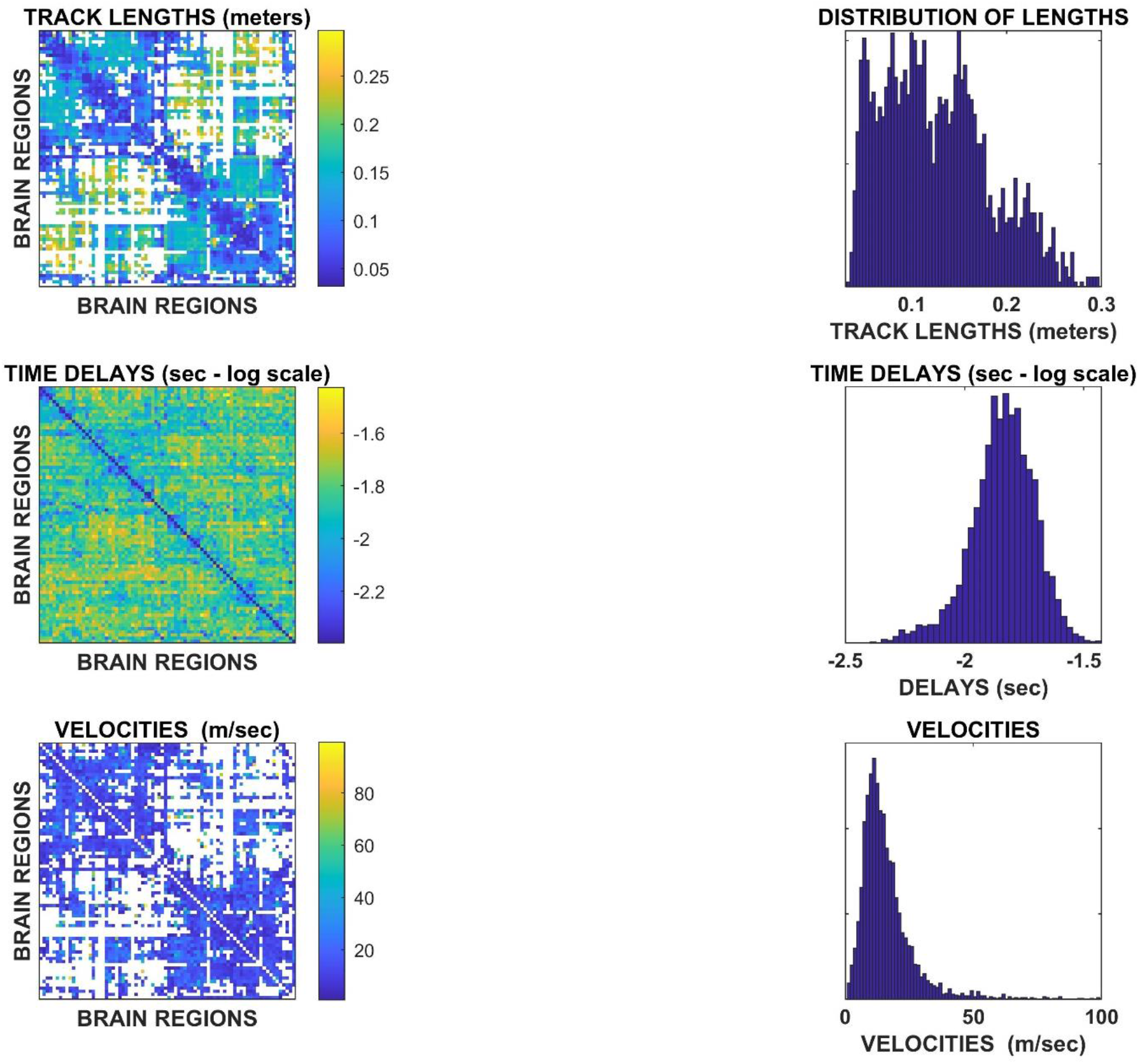
Replication of the main results on an independent cohort of 47 young healthy subjects, based on source-reconstructed MEG and tractography (parcellation according to the AAL atlas is shown).

**Fig. S14.**
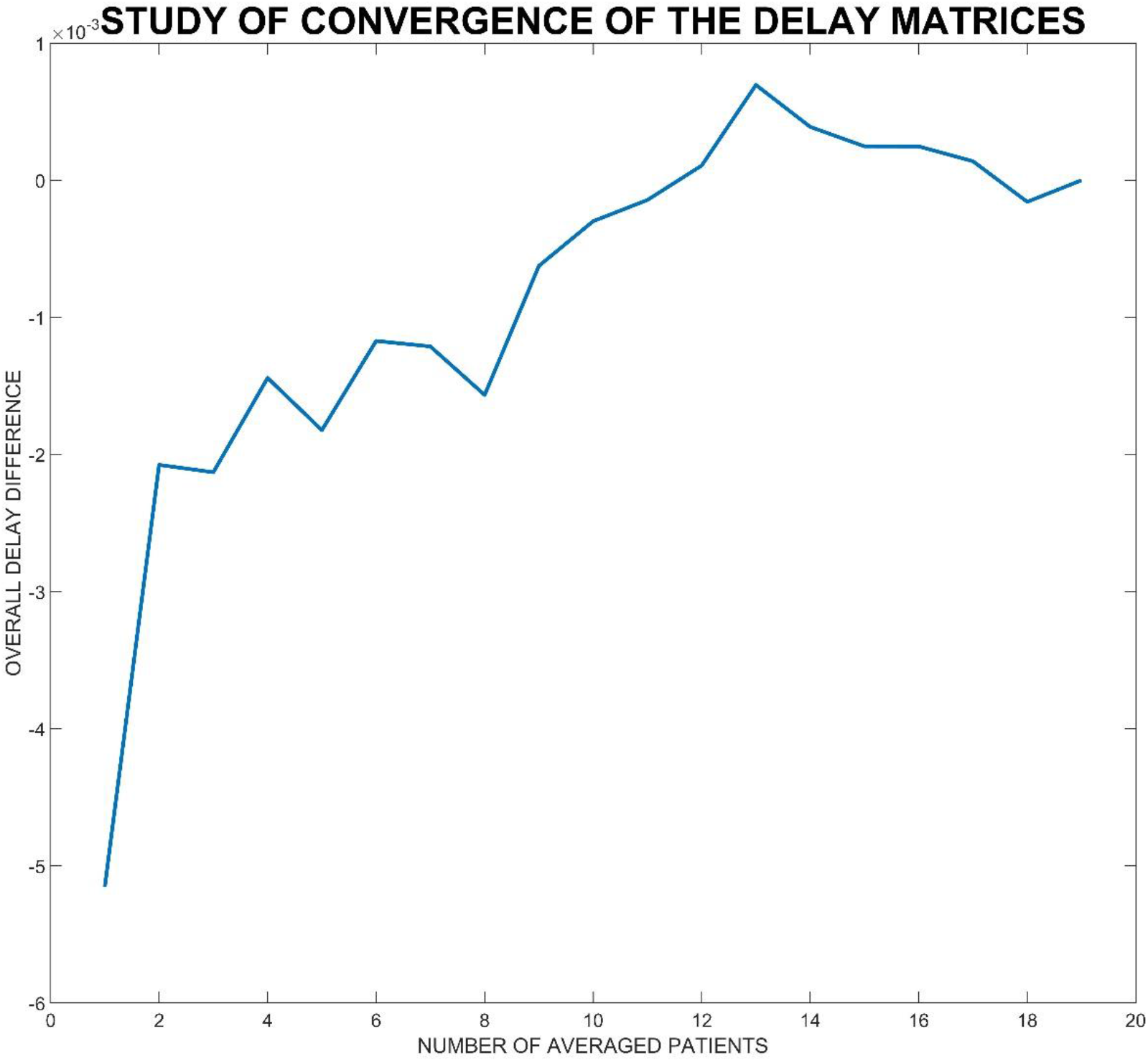
Analysis of convergence. On the x axis, the image shows the number of patients over which the delays have been averaged. The y axis shows the mean difference between the average delays (i.e., across all edges) estimated in x subjects and the average delays estimated in x-1 subjects. The differences converge around zero as the delays were averaged across a growing number of participants.

